# Transcription factor ScWRKY4 in sugarcane negatively regulates the resistance to pathogen infection through the JA signaling pathway

**DOI:** 10.1101/2023.07.04.547636

**Authors:** Dongjiao Wang, Wei Wang, Shoujian Zang, Liqian Qin, Yanlan Liang, Peixia Lin, Yachun Su, Youxiong Que

## Abstract

WRKY transcription factor, the transcriptional regulators unique to plants, plays an important role in plant defense response to pathogen infection. However, the disease resistance mechanism of *WRKY* gene in sugarcane remains unclear. Previously, we identified a *ScWRKY4* gene, a member of class IIc of the WRKY gene family, from the sugarcane cultivar ROC22. This gene could be induced by the stresses of salicylic acid (SA) and methyl jasmonate (MeJA). Interestingly, the expression of the *ScWRKY4* gene was down-regulated in smut-resistant sugarcane cultivars but up-regulated in smut-susceptible sugarcane cultivars under *Sporisorium scitamineum* stress. Besides, stable overexpression of the *ScWRKY4* gene in *Nicotiana benthamiana* enhanced susceptibility to *Fusarium solani* var. *coeruleum* and caused the down-regulated expression of immune marker-related genes. Furthermore, transcriptome analysis indicated that, the expression of most *JAZ* genes was suppressed in plant signal transduction pathway. In addition, ScWRKY4 could interact with ScJAZ13 and repress the expression of ScJAZ13. We thus hypothesized that the *ScWRKY4* gene was involved in the regulatory network of plant disease resistance, most probably through the JA signaling pathway. The present study depicted the molecular mechanism of the *ScWRKY4* gene involved in sugarcane disease resistance and laid the foundation for the subsequent investigation.

**Highlight:** Transgenic plants overexpressing the *ScWRKY4* gene negatively regulated resistance to pathogen by inhibiting the expression of the *JAZ* genes.

## Introduction

Sugarcane (*Saccharum* spp.) is a cash crop grown mainly in subtropical and tropical regions and plays an important role in the agricultural economy of China (Singels *et al*., 2021). Like other crops, biotic and abiotic stresses, such as drought, low temperature and pathogenic fungi, are the main factors restricting sugarcane production (Rajput *et al*., 2021). Sugarcane smut, which seriously endangers the healthy and sustainable development of sugarcane industry, is a worldwide fungal disease caused by the fungus *Sporisorium scitamineum* (Bhuiyan *et al*., 2021).

When plants encounter stress during growth and development, they will form a complex defense regulatory network, and transcription factors perform a critical function during this period (Singh *et al*., 2002). WRKY transcription factors, one of the largest families of transcription factors in plants, are widely involved in plant responses to biotic, abiotic and hormonal stresses (Jiang *et al*., 2017), especially in the formation of plant disease resistance (Rushton *et al*., 2010). They have a 60 amino acid long DNA binding domain, which is characterized by a highly conserved WRKYGQK core motif at the N-terminal and a CX_4–5_CX_22–23_HXH zinc finger motif at the C-terminal (Bakshi and Oelmüller, 2014). According to the number of typical WRKY domains and the type of zinc finger motif, WRKY family members can be divided into three groups, group I has two WRKY domains, while group II or III has only one WRKY domain (Eulgem *et al*., 2000). C_2_H_2_ (CX_4-5_CX_22-23_HX_1_H), where X can be any amino acid, is the zinc finger structure possessed by members of groups I and II, however it is C_2_HC (CX_7_CX_23_HXC) in group III (Eulgem *et al*., 2000). Based on the homology, the members of group II WRKY family can be further classified as five subgroups, IIa, IIb, IIc, IId and IIe (Zhang and Wang, 2005).

Plants have evolved a complex mechanism to respond to a series of biotic stresses (Gull *et al*., 2019). The regulatory role of WRKY transcription factors in plant immune response to various biotic stresses has been extensively studied (Wani *et al*., 2021), with functions including positive and negative regulation in plant defense responses to various pathogen infection (Pandey and Somssich, 2009). In *Vitis vinifera*, transient silencing of *VqWRKY31* reduced resistance to powdery mildew (Yin *et al*., 2022). In *Juglans regia*, silencing of *JrWRKY21* significantly reduced the resistance of walnuts to *Colletotrichum gloeosporioides*, while the disease resistance was significantly enhanced in walnut overexpressing *JrWRKY21* (Zhou *et al*., 2022). In *Oryza sativa*, knock out mutations of the OsWRKY53 transcription factor thickened sclerenchyma cell walls, thereby conferring higher resistance to rice bacterial blight (Xie *et al*., 2021). In *Triticum aestivum*, TaWRKY19 negatively regulated stripe rust resistance through the production of reactive oxygen species (ROS) and transcriptional repression of *TaNOX10* (Wang *et al*., 2022b). Notably, when subjected to biotic stresses, WRKY transcription factors could activate the expression levels of genes related to salicylic acid (SA) and jasmonic acid (JA) signaling pathways in plants, and then responded to different biotic stresses (Jiao *et al*., 2018). For example, AtWRKY70, an important member of the SA- and JA-regulated defense signaling pathway, was induced by SA and repressed by JA, thereby activating SA-inducible genes and repressing JA-responsive genes that were further involved in defense responses (Li *et al*., 2004). Previous research also demonstrated that overexpression of *AtWRKY70* increased the resistance of Arabidopsis to *Pseudomonas syringae* pv *tomato* (Li *et al*., 2004). Nevertheless, GhWRKY70 negatively regulated tolerance to *Verticillium dahliae* in *Gossypium hirsutum* by up-regulating the expression of SA-related genes and repressing the expression of JA-related genes (Xiong *et al*., 2019). In sugarcane, *Sc-WRKY* gene (GeneBank accession number GQ246458) belongs to class IIc was first cloned and its expression could be induced by the stress of both *S. scitamineum* and SA (Liu, 2012). In addition, the expression of sugarcane class IIc *ScWRKY3* gene (GeneBank accession number: MK034706) was up-regulated in smut-susceptible variety ROC22, but remained unchanged in smut-resistant variety Yacheng05-179, and its expression was inhibited by SA and MeJA treatments (Wang *et al*., 2018b). Li et al. identified 154 members of the SsWRKY gene family in the *Saccharum spontaneum* genome, and RNA-seq analysis revealed that *SsWRKYs* displayed different temporal and spatial expression patterns in different developmental stages, of which, 52 *SsWRKY* genes were expressed in all tissues, four *SsWRKY* genes were not expressed in any tissues, 21 *SsWRKY* genes may be involved in photosynthesis (Li *et al*., 2020). Besides, Javed et al. described 53 *ShWRKY* genes in *Saccharum* spp. hybrid R570, of which four genes, *ShWRKY13-2/39-1/49-3/125-3*, were significantly up-regulated in sugarcane cultivars LCP85-384 resistant to leaf scald (Javed *et al*., 2022). From all the above, the disease resistance regulatory mechanisms of *WRKY* genes had been widely reported in other species, however there were only few and superficial reports on the disease resistance functions of sugarcane *WRKY* genes.

We have previously reported that, the expression of *ScWRKY4* gene, which belonged to class IIc of WRKY family, could be induced under SA, methyl jasmonate (MeJA) and smut stress (Wang *et al*., 2018a). What interests us most is that, the expression of the *ScWRKY4* gene was down-regulated in smut-resistant sugarcane cultivars but up-regulated in smut-susceptible sugarcane cultivars under *S. scitamineum* stress. In the present study, we also found both transient and stable overexpression of the *ScWRKY4* gene in *Nicotiana benthamiana* enhanced the susceptibility of tobacco plants to *Fusarium solani* var. *coeruleum* and caused down-regulated expression of immune marker genes. Strikingly, after inoculation with pathogen, the expression of most JAZ-related genes was repressed in transgenic tobacco plants stably overexpressing *ScWRKY4* gene. Further experiments indicated that, ScWRKY4 interacted with ScJAZ13 and could inhibit the expression of ScJAZ13. Our study aimed to explore the regulatory network/mechanism of disease resistance for the *ScWRKY4* gene and provided a theoretical basis for subsequent studies on members of the WRKY gene family in sugarcane.

## Materials and methods

### Plant materials, culture conditions and pathogen inoculation

The two smut-resistant sugarcane cultivars (YZ01-1413 and YT96-86), and two smut-susceptible sugarcane cultivars (YZ03-103 and FN39) used in this study were provided by the Key Laboratory of Sugarcane Biology and Genetic Breeding, Ministry of Agriculture and Rural Affairs (Fuzhou, China). The above cultivars were inoculated with *S. scitamineum* according to the methods of Que et al. (Que *et al*., 2014). Three single buds were collected at 0, 1, 3 and 7 days post inoculation (dpi), and then stored in -80°C refrigerator after freezing in liquid nitrogen.

The gateway primers (Supplementary Table S1) were designed to construct the *ScWRKY4* gene into the overexpression vector pEarleyGate 203. The empty pEarleyGate 203 (*35S::00*) and the fusion vector pEarleyGate 203-*ScWRKY4* (*35S::ScWRKY4*) were transiently overexpressed in *N. benthamiana* by the *Agrobacterium*-mediated method. After 1 d, two leaves were taken from each plant and used for real-time quantitative PCR (RT-qPCR) and 3,3’-diaminobenzidine (DAB) histochemical analysis, respectively (Wang *et al*., 2020). The fungal pathogen *F. solani* var. *coeruleum* was inoculated into the over expressed *N. benthamiana* leaves for 1 day, and then the symptoms (phenotype and DAB) of *N. benthamiana* leaves were observed and the relative transcription levels of tobacco immune related marker genes were calculated (Wang *et al*., 2020).

*Agrobacterium* tumefaciens carrying pEarleyGate 203-*ScWRKY4* was stably transformed into *N. benthamiana* by leaf-disc method (Müller *et al*., 1987). The T_3_ generation of plants overexpressing *ScWRKY4* gene was screened on MS (murashige and skoog) medium supplemented with glufosinate. When the plants grew to 5–8 leaves old, the DNA of plant leaf was extracted and diluted to 25 ng/L. The pEarleyGate203-*ScWRKY4* plasmid was used as a positive control and wild-type (WT) *N. benthamiana* as a negative control. Genomic DNA of transgenic plants was used as a template for PCR amplification and electrophoresis detection (Su *et al*., 2020). RNA was extracted from the leaves of the plants, and the expression of *ScWRKY4* gene in the transgenic plants was measured using WT plants as the control (Sun *et al*., 2023). Transgenic plants overexpressing the *ScWRKY4* gene and WT were inoculated with the *F. solani* var. *coeruleum* and the phenotypic leaves were analyzed by DAB and trypan blue staining (Su *et al*., 2022). Samples were taken at 0 and 2 dpi for subsequent detection of immune marker related genes, determination of physiological indicators, and transcriptome analysis.

### RNA extraction and RT-qPCR analysis

The total RNA was extracted by TRIzol method (Connolly *et al*., 2006). Refer to the kit instructions of Hifair® III 1st Strand cDNA Synthesis SuperMix for qPCR (gDNA digester plus), RNA was reverse transcribed from extracted treated material into first strand cDNA, which was used as a template for quantitative detection of target gene expression levels. The obtained RNA and cDNA were tested for quality by 1.0% agarose gel electrophoresis. Primer Premier 5 software was used to design specific quantitative PCR primers (primer pair: *ScWRKY4*-Q-F/R) for the *ScWRKY4* gene, using glyceraldehyde-3-phosphate dehydrogenase (GAPDH) (GenBank Accession Number. CA254672) as the internal reference gene (primer pair: *GAPDH*-Q-F/R) (Supplementary Table S1) (Que *et al*., 2009). Ten tobacco immune-related marker genes including hypersensitive response (HR) marker genes *NbHSR201, NbHSR203* and *NbHSR515*, ethylene (ET) synthesis-dependent genes *NbACO-like* and *NbACO*, SA signaling pathway-related genes *NbPR2* and *NbPR3*, JA signaling pathway-related genes *NbLOX1* and *NbDEF1*, and ROS related genes *NbCAT1* and *NbGST1* (Supplementary Table S1) were selected and their expression in tobacco leaves was examined (Sun *et al*., 2023). The ABI 7500 Real-time PCR System (USA) was used for RT-qPCR detection, and the quantitative reaction system was prepared and the program was set up according to the SYBR Green PCR Master Mix Kit instructions. Three technical replicates were set up for each sample, and negative controls were used as templates with sterile water. The relative gene expression was calculated using 2^-ΔΔCT^ (Livak and Schmittgen, 2001), and the significance level of the experimental data was analyzed using DPS 9.50 software and histograms were plotted using Origin 2022 software.

### Gene ontology (GO) enrichment analysis of ShWRKY gene family members

GO annotate our previously identified ShWRKY gene family members (Wang *et al*., 2022a) based on the eggNOG-mapper v2 database (http://eggnog-mapper.embl.de/) with parameters set to default parameters (Cantalapiedra *et al*., 2021). The obtained data were then analyzed by GO enrichment using Tbtools software (Chen *et al*., 2020) and visualized by the online software chiplot (https://www.chiplot.online/).

### Bioinformatics analysis of ScWRKY4

Structural domain prediction of the ScWRKY4 protein was conducted using the SMART website (https://smart.embl.de/). Protein sequence alignment was performed using MUSCLE v3.7 (Edgar, 2004), and the phylogenetic evolutionary tree of ShWRKY proteins and ScWRKY4 protein was constructed using IQ-TREE in PhyloSuite software, and the replicate value was set to 1000 times (Zhang *et al*., 2020). Beautify the evolutionary tree with the help of EvolView online website (https://evolgenius.info//evolview-v2) (Subramanian *et al*., 2019). The conserved motif information of *ScWRKY4* gene was obtained through the analysis of MEME online website (http://meme-suite.org/index.html), and the parameter settings showed 10 conserved motifs and the rest with default parameters (Bailey *et al*., 2009). Multiple sequence alignment of the *ScWRKY4* gene with the *ShWRKY* genes was performed by DNAMAN software to obtain exon-intron structure information of the *ScWRKY4* homologous gene from the *S*. spp hybrid R570 genome (Garsmeur *et al*., 2018) GFF3 file and gene structure of the *ScWRKY4* homologous gene were visualized using TBtools software (Chen *et al*., 2020). Retrieved 2000 bp promoter sequences upstream of homologous genes were retrieved and *cis*-acting regulatory elements in promoter sequences were predicted using PlantCARE online website (https://bioinformatics.psb.ugent.be/webtools/plantcare/html/) (Lescot *et al*., 2002).

### Hormone content and enzyme activity determination

The contents of endogenous hormones SA and JA were determined in leaves of WT and transgenic tobacco plants at 0 dpi and 2 dpi respectively, according to the instructions of the plant SA ELISA kit and JA ELISA kit (Enzyme-linked Biotechnology, Shanghai, China). Similarly, the enzymatic activities of catalase (CAT) and glutathione S transferases (GST) were measured in leaves of WT and transgenic tobacco plants at 0 dpi and 2 dpi respectively, according to the CAT ELISA kit and GST ELISA kit (Enzyme-linked Biotechnology, Shanghai, China).

### Transcriptome data analysis

RNA samples from WT plants inoculated with *F. solani* var. *coeruleum* at 0 d and 2 d were named WT-CK and WT-T, respectively. And RNA samples from *ScWRKY4-*overexpressed transgenic tobacco plants inoculated with *F. solani* var. *coeruleum* at 0 d and 2 d were named *ScWRKY4*-CK and *ScWRKY4*-T, respectively. Three biological replicates were set up for each of the above samples, with a total of 12 samples (WT-CK1, WT-CK2, WT-CK3, WT-T1, WT-T2, WT-T3, *ScWRKY4*-CK1, *ScWRKY4*-CK2, *ScWRKY4*-CK3, *ScWRKY4*-T1, *ScWRKY4*-T2 and *ScWRKY4*-T3). RNA-Seq sequencing was entrusted to Genedenovo. The raw reads were quality controlled using fastp to filter low quality data and get clean reads (Chen *et al*., 2018). A comparative analysis based on the *N. benthamiana* (https://sefapps02.qut.edu.au/benWeb/subpages/downloads.php) database was carried out and annotated using the HISAT2 software (Langmead and Salzberg, 2012). Bioinformatic analysis was performed using Omicsmart (http://www.omicsmart.com), a dynamic real-time interactive online platform for data analysis.

### Yeast two-hybrid

The recombinant plasmids pGBKT7-ScWRKY4 and pGADT7-ScJAZs (JAZ6, 8, 9, 10, 11, 13 and 14) constructed earlier by our group were used in the yeast two-hybrid system. The positive control pGADT7-T+pGBKT7-p53, negative control pGADT7-T+pGBKT7-lam and combination of plasmid pGBKT7-ScWRKY4+pGADT7-ScJAZs were co-transformed into Y2HGold yeast strain, respectively. The transformation method was carried out according to the instructions of Y2HGold Chemically Competent Cell. After transformation, the Y2HGold strain containing the above plasmid combination was spread on tryptophan -leucine (SD/-Trp-Leu) deficient medium plates, and cultured in a 28°C incubator for 2–3 d. Single colony was selected and culture in SD/-Trp-Leu liquid medium for about 18 h, and then diluted four gradients (10^-1^, 10^-2^, 10^-3^, 10^-4^) with sterile water, with 10 µL to spotted on SD/-Trp-Leu and tryptophan-leucine-histidine-adenine (SD/-Trp-Leu-His-Ade) deficient medium plates, respectively. After that, it was cultivated in a 28°C incubator for 2–3 d until the yeast colonies grew, and the growth of the yeast colonies was recorded by taking pictures. The interaction of proteins in the yeast was judged according to whether the yeast colonies could grow normally.

### Bimolecular fluorescent complimentary (BiFC)

Using the gateway method to construct *ScWRKY4* gene and *ScJAZ13* gene into pEarlyGate201 and pEarlyGate202 vectors, respectively. The BiFC recombinant vectors pEarlyGate201-*ScWRKY4*, pEarlyGat202-*ScJAZ13*, the empty-loaded pEarlyGate201-YN and pEarlyGate202-YC were transformed into *Agrobacterium* GV3101 strains, respectively, according to the principle of equal proportion pairing. Injected into 5–8 leaf-aged *N. benthamiana* leaves with the same growth, and after culturing for 3 d at room temperature, about 1 cm^2^ of tobacco leaves were cut to make glass slides, and the specimens were examined under a Leica TCS SP8 laser confocal microscope (Leica Microsystems (Shanghai) Trading Co., Ltd., Mannheim, Germany), and the yellow fluorescent protein (YFP) was observed with a 10×lens and a YFP filter (561 nm excitation wavelength).

### Detection of protein inhibition using green fluorescent protein (GFP)

The *ScJAZ13* gene was constructed into the vector pCAMBIA2300 with GFP tag, and the fusion vector pCAMBIA2300-ScJAZ13 was then transformed into *Agrobacterium* tumefaciens strain GV3101. *Agrobacterium* containing the target fragment and H2B-mCherry (Howe *et al*., 2012) were subsequently collected and diluted with MS blank medium and mixed in equal proportions to OD = 0.8. The 1.0 mL bacterial solution (containing 200 μmol-L^-1^ acetosyringone) was injected with a sterile syringe into 5–8 leaf old WT plants and transgenic plants overexpressing *ScWRKY4* gene, respectively. After incubation for 2 d at 28°C under light for 16 h/dark for 8 h, the leaves were collected and the GFP was observed under a Leica TCS SP8 laser confocal microscope (Leica Microsystems (Shanghai) Trading Co., Ltd., Mannheim, Germany) using a 10×lens, 488 nm green fluorescence excitation wavelength.

## Results

### GO annotation of ShWRKY gene family

In previous study, we identified 60 *ShWRKY* genes from the genome of *S*. spp. hybrid cultivar R570 (Wang *et al*., 2022a). These *ShWRKY* genes were constitutively expressed in sugarcane tissues and their expression could be induced by the smut pathogen (Wang *et al*., 2022a). To further explore the function of *ShWRKY* genes, the GO annotation analysis was performed. The results showed that *ShWRKY* genes were annotated in the biological process (38%), molecular function (36%), and cellular component (26%) categories (Fig. 1A and Supplementary Table S2). In the biological process, *ShWRKY* genes were mainly enriched in biotic stress response and defense response items (Fig. 1B and Supplementary Table S2). For molecular function, 26 *ShWRKY* genes were enriched in transcriptional regulatory activity and DNA-binding transcription factor activity items (Fig. 1C and Supplementary Table S2). While in the cellular component, most of the *ShWRKY* genes were enriched in the nucleus, the cellular component items (Fig. 1D and Supplementary Table S2).

**Fig. 1.**
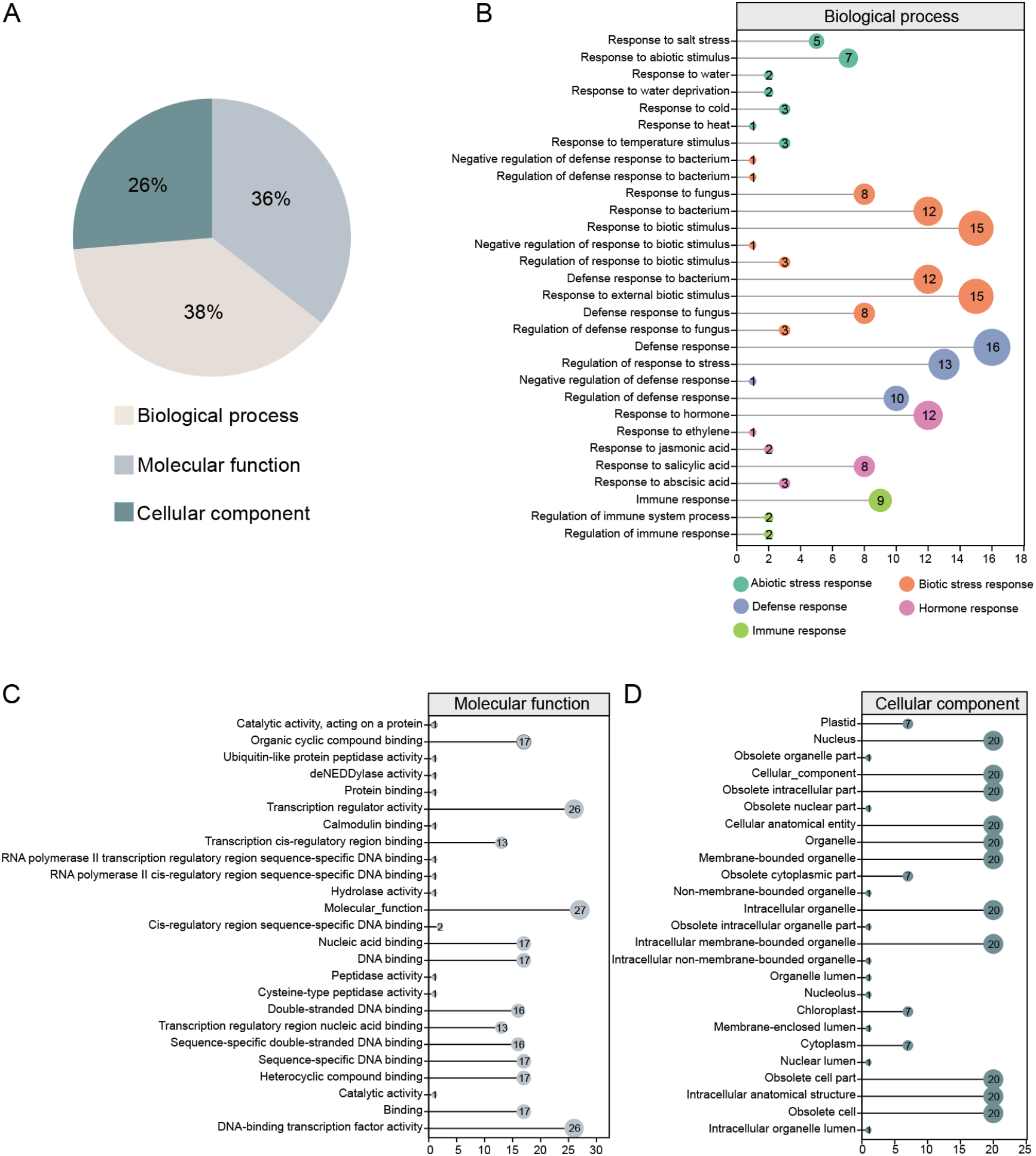
GO annotation of ShWRKY gene family in *Saccharum*. spp. hybrid cultivar R570. (A) The category of GO function. (B) GO function of ShWRKY gene family in molecular function. (C) GO function of ShWRKY gene family in biological process. (D) GO function of ShWRKY gene family in cellular component.

### Characteristics of the ScWRKY4 in sugarcane

Using the leaf cDNA of sugarcane ROC22 as a template, the *ScWRKY4* gene was cloned. The ORF (137–877 bp) of this gene was 741 bp in length, encoding 246 amino acids, and with a typical WRKY structural domain (Fig. 2A). A phylogenetic tree of ScWRKY4 protein was constructed using the ShWRKY proteins as a reference, and the structure showed that ScWRKY4 protein belongs to subgroup IIc (Fig. 2B). The results of multiple sequence comparison indicated that ScWRKY4 had the highest similarity of 99.19% with ShWRKY36 (Supplementary Table S3), suggesting their functional similarity. A total of 10 conserved motifs of ScWRKY4 protein were observed using MEME software. The ScWRKY4 protein contained only Motif 1, Motif 2, Motif 3, and Motif 6 (Fig. 2C). Gene structure analysis revealed that the *ScWRKY4* gene had four exons and three introns (Fig. 2D). Prediction of promoter element in the first 2000 bp upstream of *ScWRKY4* gene suggested that several *cis*-acting regulatory elements were associated with stress, growth and hormone response (Fig. 2E and Supplementary Table S4). Of these, one stress-related *cis*-acting regulatory element LTR, was involved in the low temperature response. Two growth and development related *cis*-acting regulatory elements, such as the ARE, was essential for anaerobic induction, and the RY-element acted as a component involved in seed specificity. There were also four hormonal response related *cis*-acting regulatory elements, included MeJA-responsive element TGACG-motif, auxin-responsive element TGA-element, ABA-responsive element ABRE and gibberellin-responsive element P-box (Fig. 2E and Supplementary Table S4). It is thus hypothesized that *ScWRKY4* gene may be involved in sugarcane growth and development and in the defence process against biotic stresses.

**Fig. 2.**
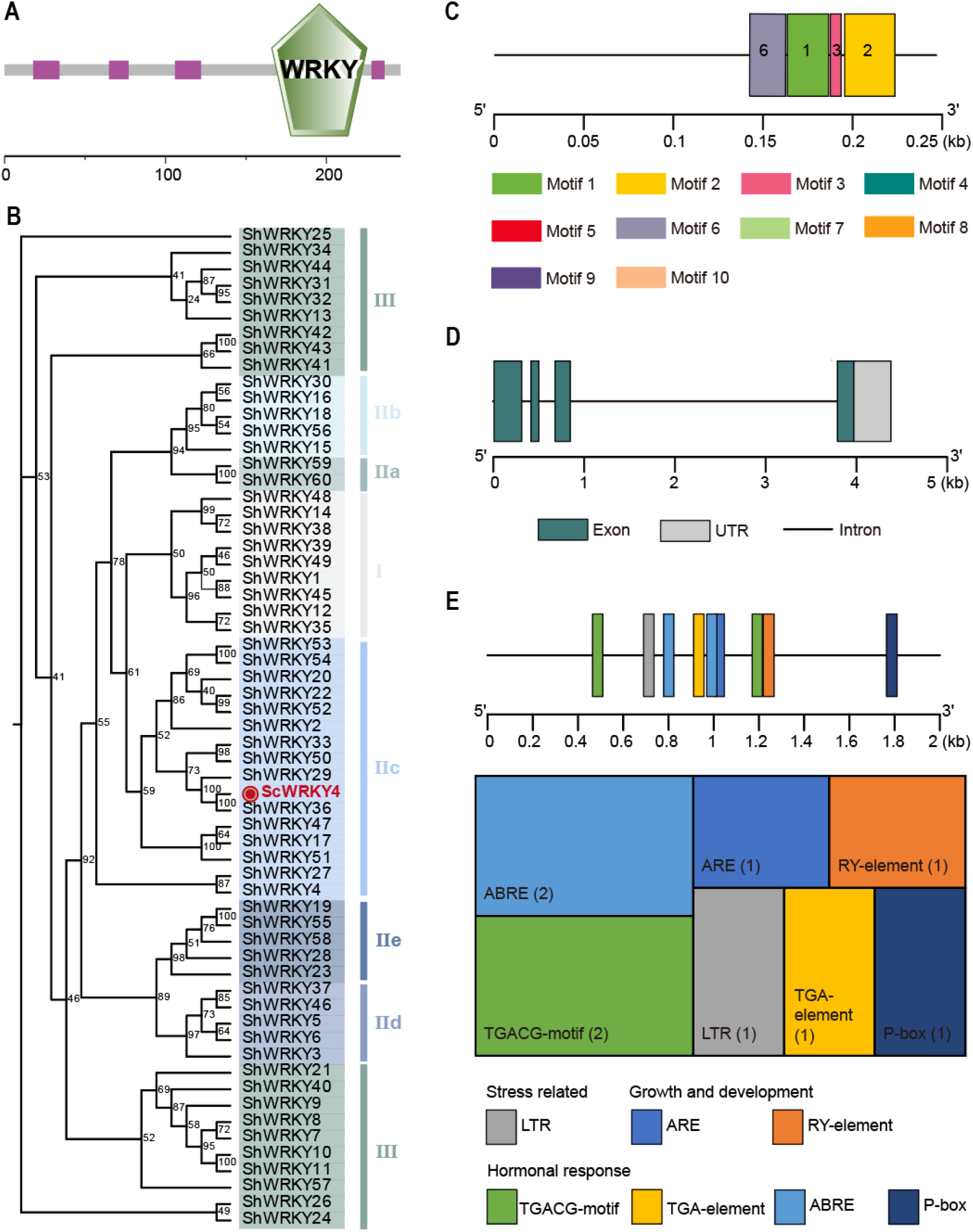
Characterization of ScWRKY4 in sugarcane. (A) Conserved domains of ScWRKY4 protein. (B) A phylogenetic tree of ScWRKY4 protein. ScWRKY4 protein was shown in red bold font. (C) Conserved motifs of ScWRKY4 protein. (D) Structure of *ScWRKY4* gene. (E) Distribution and functional prediction of *cis*-acting regulatory elements of *ScWRKY4* gene promoter. The numbers in parentheses in the figure represented the number of *cis-*acting regulatory element.

### The expression profile of ScWRKY4 gene

Previous study showed that, the expression level of *ScWRKY4* gene reached a peak value at 12 h under both SA and MeJA treatments, which was 2.69-folds and 2.38-folds that of the control group, respectively (Fig. 3A). Fig. 3A demonstrated the expression level of *ScWRKY4* gene in four different sugarcane genotypes interacting with smut pathogen. As a result, the *ScWRKY4* gene was down-regulated in two smut-resistant sugarcane cultivars (YZ01-1413 and YT96-86), while up-regulated in two smut-susceptible sugarcane cultivars (YZ03-103 and FN39) at 7 dpi. Therefore, it is speculated that the *ScWRKY4* gene may negatively regulate the defence of sugarcane against smut fungus infection.

**Fig. 3.**
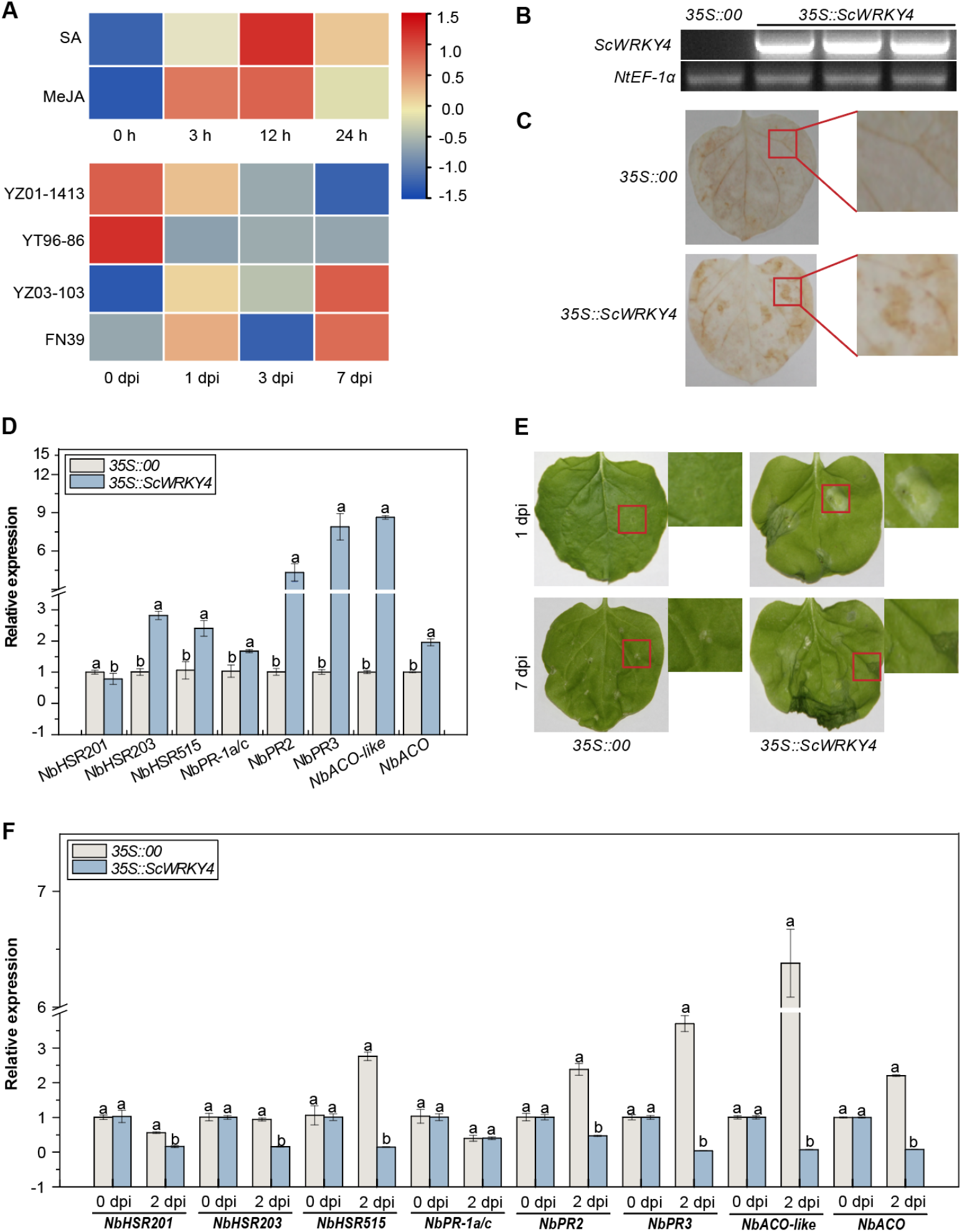
The function of *ScWRKY4* gene. (A) Relative expression of *ScWRKY4* genes in sugarcane under SA, MeJA and smut stress (YZ01-1413 and YT96-86 were smut-resistant sugarcane cultivars, YZ03-103 and FN39 were smut-susceptible sugarcane cultivars). (B) Semi-quantitative PCR amplification of *ScWRKY4* in *N. benthamiana* leaves. *35S::00*, the empty vector pEarleyGate 203. *35S::ScWRKY4*, pEarleyGate 203-*ScWRKY4*. (C) DAB staining. (D) RT-qPCR expression of eight immunity-associated marker genes in the *N. benthamiana* leaves. (E) Phenotype of *N. benthamiana* leaves after inoculation with *F. solani var. coeruleum*. (F) The transcripts of immunity-associated marker genes in the *N. benthamiana* leaves after inoculation with *F. solani* var. *coeruleum* at 2 dpi. Data were normalized to the *NbEF-1α* expression level. All data points were means ± standard errors (*n* = 3). Different letters on the columns represented significant differences calculated by Duncan’s new multiple range test (*P*<0.05).

Next, we transiently transformed *ScWRKY4* gene into *N*. *benthamiana* leaves by *Agrobacterium* mediated method, and the ScWRKY4 protein was successfully detected in tobacco leaves at 1 d (Fig. 3B). DAB staining revealed, that tobacco leaves transiently overexpressing the *ScWRKY4* gene (*35S::ScWRKY4*) had darker browning and a more pronounced allergic response than the control group (*35S::00*) (Fig. 3C). The expression of seven HR marker gene was significantly up-regulated except for *NbHSR201* (Fig. 3D). After inoculation with *F. solani* var. *coeruleum* for 7 d, *35S::ScWRKY4* tobacco leaves began to wilting, and the degree was more serious than that of the *35S::00* (Fig. 3E). Compared with the *35S::00* plants, in the *35S::ScWRKY4* tobacco leaves at 2 dpi, the expression of seven immune related genes were down regulated, except that the expression of *NbPR-1a/c* did not change (Fig. 3F). The results suggest that ScWRKY4 may be a negatively regulated transcription factor.

### Stable overexpression of ScWRKY4 negatively regulates resistance to pathogen infection

To further examine the disease resistance of *ScWRKY4* gene, we genetically transformed it into *N. benthamiana* and cultured it to T_3_ generation by screening on MS plates supplemented with herbicide, and finally obtained the transgenic positive lines of *ScWRKY4* (OE) (Supplementary Fig. S1A). A target band consistent with the size of the overexpression vector plasmid was identified in all transgenic plants, indicating that the *ScWRKY4* gene was successfully inserted into the genome of *N. benthamiana*, which could be used for the next step of disease resistance verification (Supplementary Fig. S1B).

From the perspective of phenotype, after inoculation with *F. solani* var. *coeruleum*, the leaves of OE plants showed significant wilting and yellowing with obvious disease spot symptoms compared to the WT at 7 dpi (Fig. 4A). Compared with WT, the DAB (H_2_O_2_ accumulation) and trypan blue staining (cell death) of OE leaves were deeper. Besides, the SA and JA contents of WT and transgenic lines plants were measured at 0 dpi and 2 dpi. There was no significant difference in the content of SA in WT and OE, while the content of JA in OE was significantly higher than the control group, 1.28-folds higher than the control group at 0 dpi. At 2 dpi, the content of SA and JA in OE plants was significantly lower than that in WT plants, 0.93-folds and 0.89-folds respectively (Fig. 4B). Enzyme activity assay revealed that the CAT and GSTs activities of the OE plants were significantly higher than the control before inoculation, 2.44- and 1.16-folds higher than the control, respectively. However, the activities of both enzymes decreased to the control level at 2 dpi (Fig. 4C). Finally, when the expression of immune marker-related genes was examined at 0 and 2dpi, the expression of the HR marker gene (*NbHSR201* and *NbHSR515*), the SA signalling pathway-related genes (*NbPR2* and *NbPR3*), the JA signalling pathway-related genes (*NbLOX1* and *NbDEF1*), and the ROS-related gene (*NbCAT1*) were all suppressed in the OE plants compared to the WT (Fig. 4D–G). The above results demonstrated that overexpression of *ScWRKY4* gene in *N. benthamiana* enhanced the susceptibility of transgenic tobacco plants to *F. solani* var. *coeruleum* infection.

**Fig. 4.**
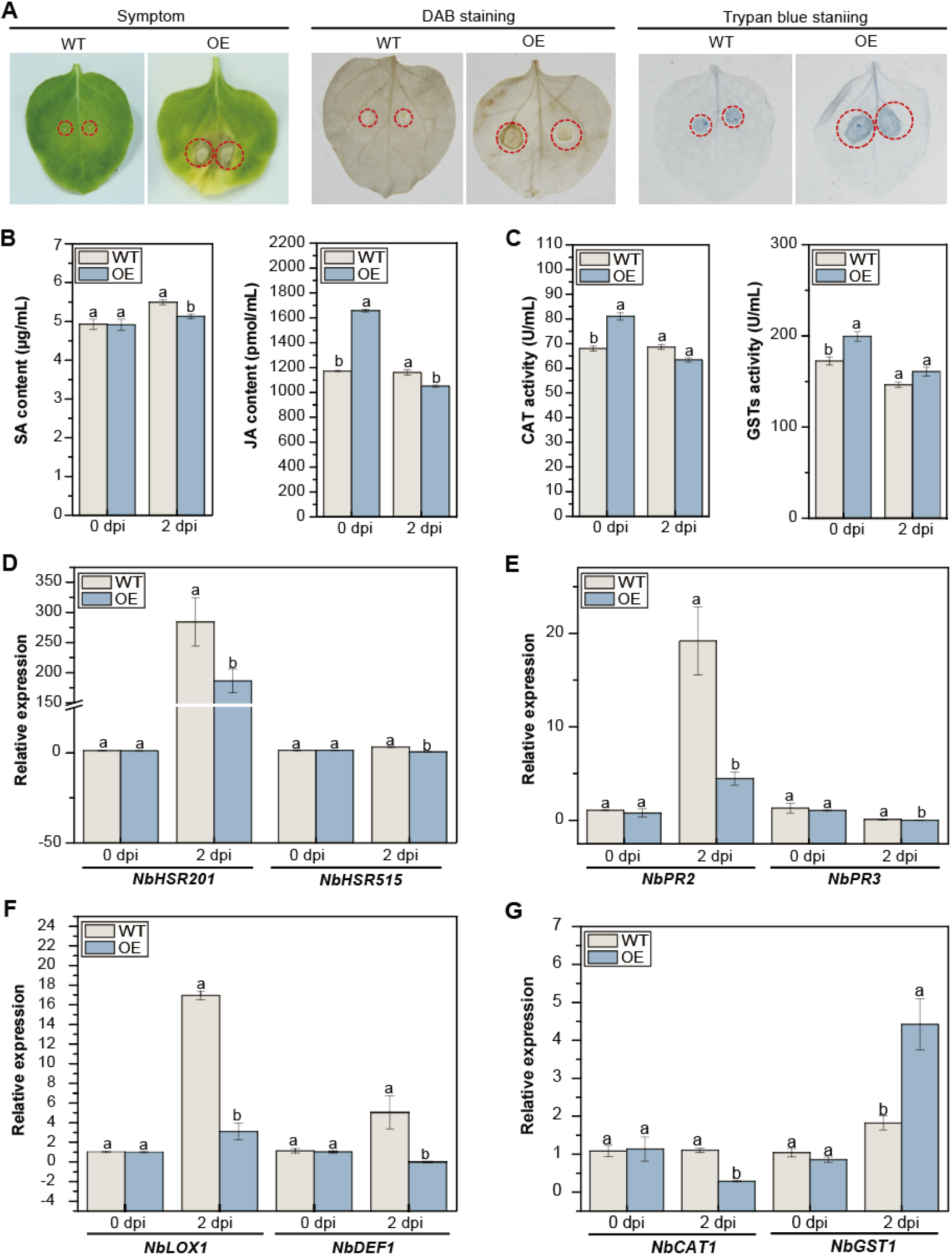
Disease resistance of *ScWRKY4* transgenic *N. benthamiana* inoculated with *F. solani* var. *coeruleum*. (A) Phenotype observation at 7 dpi. Use the red circle to mark lesion. (B) Determination of SA and JA contents in WT and OE plants at 0 dpi and 2 dpi. (C) CAT and GSTs enzyme activity assay in WT and OE plants at 0 dpi and 2 dpi. (D–G) Expression of HR marker genes, SA and JA signaling pathways, and ROS defense response related genes. WT, wild type *N. benthamiana*, OE, transgenic *N. benthamiana* plants overexpressing *ScWRKY4* gene. Data were normalized to the *NbEF-1α* expression level. All data points were means ± standard errors (*n* = 3). Different letters on the columns represented significant differences calculated by Duncan’s new multiple range test (P<0.05).

### Differentially expressed genes (DEGs) involved in disease resistance regulatory network of the ScWRKY4 gene

To obtain the DEGs involved in disease resistance, the WT and *ScWRKY4*-OE samples inoculated with *F. solani* var. *coeruleum* (WT-CK, WT-T, *ScWRKY4*-CK and *ScWRKY4*-T) at 0 d and 2 d were sequenced. The comparison combination was set as control group WT-CK-vs-WT-T and treatment group *ScWRKY4*-CK-vs-*ScWRKY4*-T. In clean reads of all samples, Q20 and Q30 were all greater than 91%, and GC content was higher than 42%, indicating that the sequencing quality was well, and the data could be used for subsequent sample analysis (Supplementary Table.S5). The regions aligned to the genome were divided into exon regions, intron regions and intergenic regions, that most of the reads could be aligned to exon regions (Supplementary Fig. S2A). The Pearson correlation coefficient between replicate samples within a group were all close to 1 (Supplementary Fig. S2B), indicating good inter-sample repeatability for the next step of between-group difference analysis.

The up-regulated DEGs in WT were significantly lower than those in OE plants when inoculated with *F. solani* var. *coeruleum* for 0 d (Fig. S3A). There were increased up-regulated DEGs in WT and decreased up-regulated DEGs in OE plants at 2 dpi (Fig. S3B). A total of 4944 up-regulated DEGs and 3903 down-regulated DEGs were identified in WT-CK-vs-WT-T, whereas a total of 8043 up-regulated DEGs and 8013 down-regulated DEGs were identified in *ScWRKY4*-CK-vs-*ScWRKY4*-T (Fig. S3C). The down-regulated DEGs in *ScWRKY4*-CK-vs-*ScWRKY4*-T was 2.05-fold higher than in WT-CK-vs-WT-T (Fig. S3C). Among the DEGs, 5488 were unique to WT-CK-vs-WT-T and 12697 to *ScWRKY4*-CK-vs-*ScWRKY4*-T (Fig. S3D). Subsequently, GO and KEGG analyses were performed to explore the disease resistance regulatory network of the *ScWRKY4* gene. We found that 10.23% of the specific DEGs in WT-CK-vs-WT-T were enriched in the response to hormone item and 10.62% of the specific DEGs were enriched in the response to endogenous stimulus item (Fig. 5A, Supplementary Table. S6). In *ScWRKY4*-CK-vs-*ScWRKY4*-T, 32.34% specific DEGs were enriched in response to stimulus item, 18.73% in response to stress item, and 6.51% in response to external biotic stimulus item (Fig. 5B, Supplementary Table. S7). Surprisingly, DEGs specific to both WT-CK-vs-WT-T and *ScWRKY4*-CK-vs-*ScWRKY4*-T were significantly enriched in the plant hormone signal transduction pathway, MAPK signaling pathway-plant pathway and plant-pathogen interaction pathway (Fig. 5C, D). By contrast, the specific DEGs in WT-CK-vs-WT-T were significantly enriched in the plant hormone signal transduction pathway (Fig. 5C), whereas the specific DEGs in *ScWRKY4*-CK-vs-*ScWRKY4*-T were significantly enriched in the plant-pathogen interaction pathway (Fig. 5D).

**Fig. 5.**
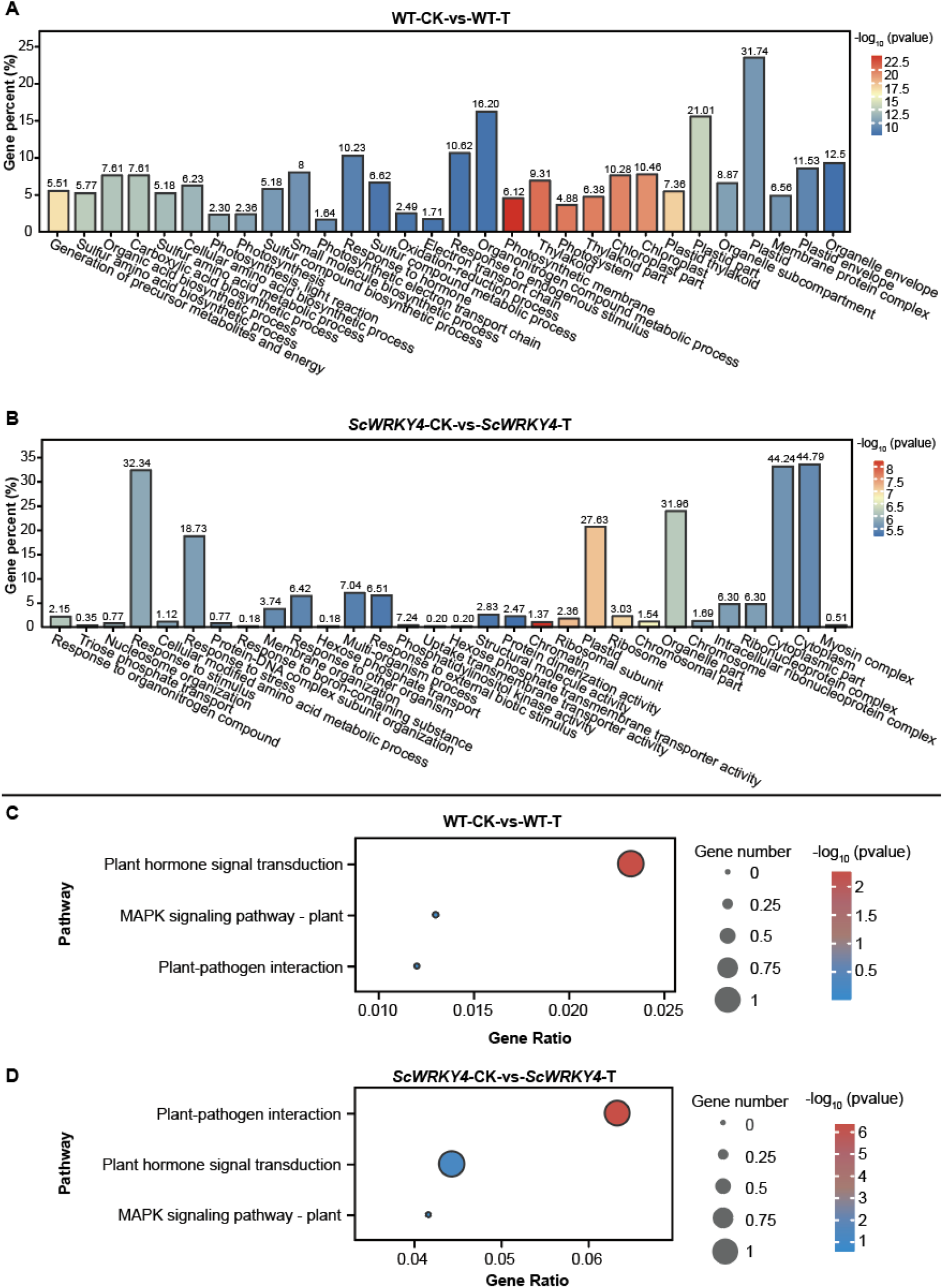
GO and KEGG enrichment of differentially expressed genes. (A) The bar graph of DEGs in the control group. (B) The bar graph of DEGs in the treatment group. (A) and (B) showed only the first 30 GO terms that were significantly enriched. (C) Bubble plot of DEGs enriched in disease resistance-related pathways in the control group. (D) Bubble plot of DEGs enriched in disease resistance-related pathways in the treatment group. Control group: WT-CK-vs-WT-T, treatment group: *ScWRKY4*-CK-vs-*ScWRKY4*-T.

### ScWRKY4 mediated the regulation of disease resistance-related pathway genes during pathogenic stress responses

To investigate the disease resistance regulatory mechanism of *ScWRKY4* gene, we further analyzed the expression of key DEGs in the disease resistance-related pathway identified above. In the plant hormone signal transduction pathway, JA and SA pathway related to stress response and disease resistance, respectively. The specific DEGs *TIFY10A* (*Nbv5tr6373992, Nbv5tr6373993, Nbv5tr6204331* and *Nbv5tr6233370*), *TIFY10B* (*Nbv5tr6348185, Nbv5tr6349145, Nbv5tr6349147, Nbv5tr6374864* and *Nbv5tr6405095*) and *TIFY11B* (*Nbv5tr6349144* and *Nbv5tr6349145*) in WT-CK-vs-WT-T were up-regulated (Fig. 6A, Supplementary Table. S8). However, the expression of these DEGs were inhibited in *ScWRKY4*-CK-vs-*ScWRKY4*-T (Fig. 6A, Supplementary Table. S8). In the plant-pathogen interaction pathway, CNGCs and CDPK were involved in the HR process of plants (Fig. 6B). CNGCs-related DEGs *CNGC1* (*Nbv5tr6233756* and *Nbv5tr6399602*) were up-regulated, and CDPK-related DEGs *CPK2* (*Nbv5tr6321646*) and *CPK5* (*Nbv5tr6397003*) were down-regulated (Fig. 6B, Supplementary Table. S8). In the MAPK signaling pathway-plant pathway, most of the DEGs in WT-CK-vs-WT-T were inhibited (Fig. 6C). Among them, the FLS2-related DEGs *XA21* (*Nbv5tr6263859*), VIP1-related DEGs *VIP1* (*Nbv5tr6405992*) and *BZIP18* (*Nbv5tr6405991*) were up-regulated and involved in defense response and late defense response for pathogen (Fig. 6C, Supplementary Table. S8). Notably, most of the DEGs in the *ScWRKY4*-CK-vs-*ScWRKY4*-T were down-regulated, among which the pathogen attack-related DEGs (*Nbv5tr6231425, Nbv5tr6339859* and *Nbv5tr6343569*) in pathogen attack were significantly down-regulated, and participated in the H_2_O_2_ production (Fig. 6C, Supplementary Table. S8).

**Fig. 6.**
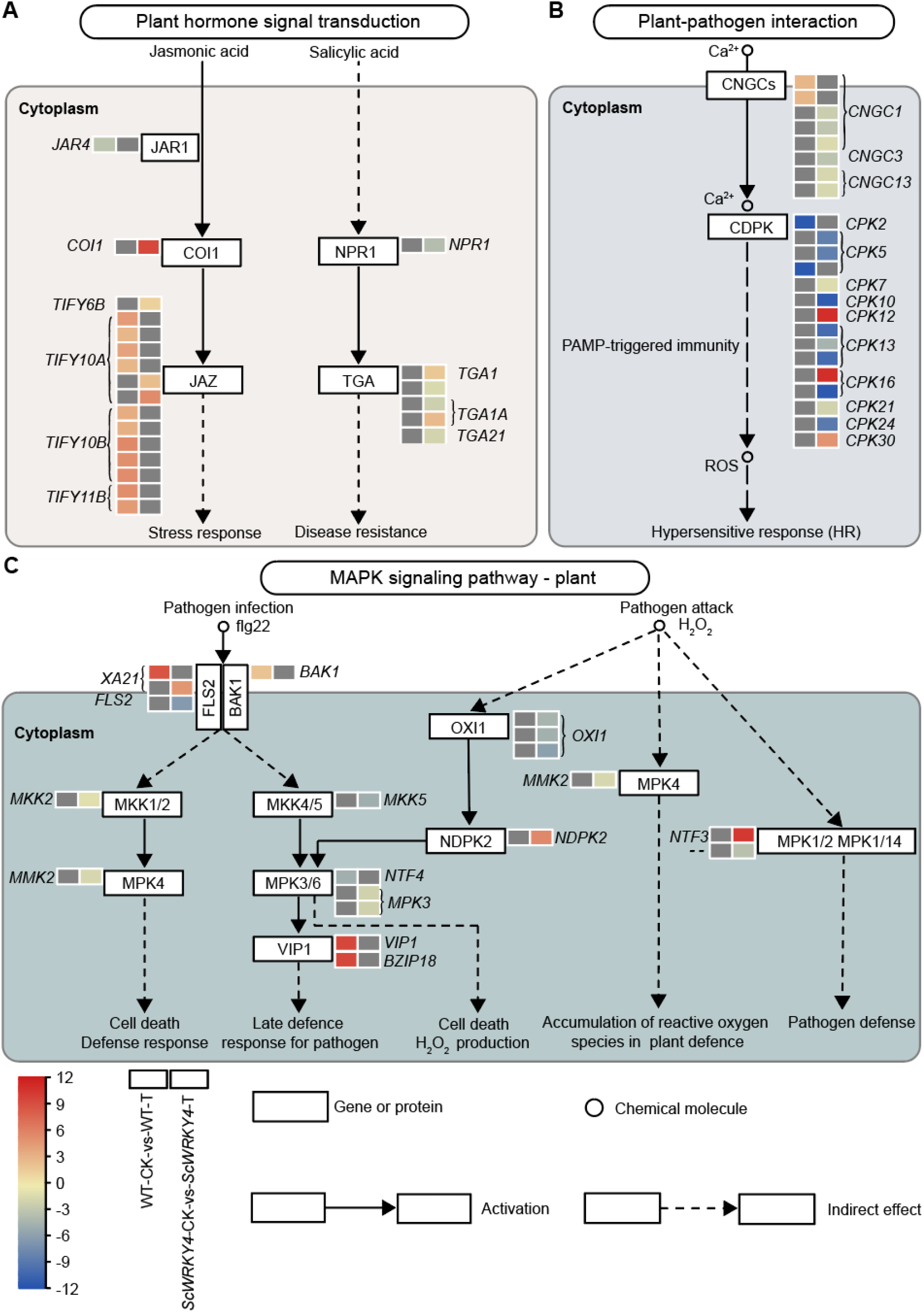
Pathways related to disease resistance and the expression levels of key genes. (A–C) The expression patterns of specific DEGs in the plant hormone signal transduction pathway, plant-pathogen interaction pathway and MAPK signaling pathway-plant pathway in the control group and the treatment group, respectively. Control group: WT-CK-vs-WT-T, treatment group: *ScWRKY4*-CK-vs-*ScWRKY4*-T. Gray box represents no expression.

### ScWRKY4 interacted with ScJAZ13 and inhibited its expression

Transcriptome analysis revealed that *JAZ* genes were mostly up-regulated in WT-CK-vs-WT-T and their expression was repressed in *ScWRKY4*-CK-vs-*ScWRKY4*-T. Therefore, we analyzed the interactions between ScWRKY4 and the seven JAZ proteins previously cloned by the group. The results of the point-to-point yeast two-hybrid indicated that all plasmid combinations grew normally on SD/-Trp-Leu medium, implying that all plasmids were successfully transformed into the yeast strain Y2HGold (Fig. 7A). However, when transferred to SD/-Trp-Leu-His-Ade medium, only the BD-ScWRKY4+AD-ScJAZ13 combination was able to grow normally, suggesting that ScWRKY4 and ScJAZ13 had a reciprocal relationship (Fig. 7A). The BiFC approach was further used to verify the interaction between ScWRKY4 and ScJAZ13. When ScWRKY4 was fused to nYFP (nYFP-ScWRKY4), and ScJAZ13 was fused to cYFP (cYFP-ScJAZ13), fluorescent complexes were formed and observed in the nucleus of *N. benthamiana* leaf cells (Fig. 7B). It is demonstrated that there is an interaction between ScWRKY4 and ScJAZ13 and that the specific protein complex is produced in the nucleus. After that, we constructed ScJAZ13 into GFP tagged vector (ScJAZ13-GFP) and transiently overexpressed it in the leaves of WT and OE plants, respectively. Two days later, the fluorescence intensity of ScJAZ13-GFP could be seen to be significantly higher in the leaves of WT plants than in the leaves of OE plants (Fig. 7C). Consequently, it is hypothesized that ScWRKY4 may repress the expression of ScJAZ13 protein and thus negatively regulated the disease resistance to pathogen infection.

**Fig. 7.**
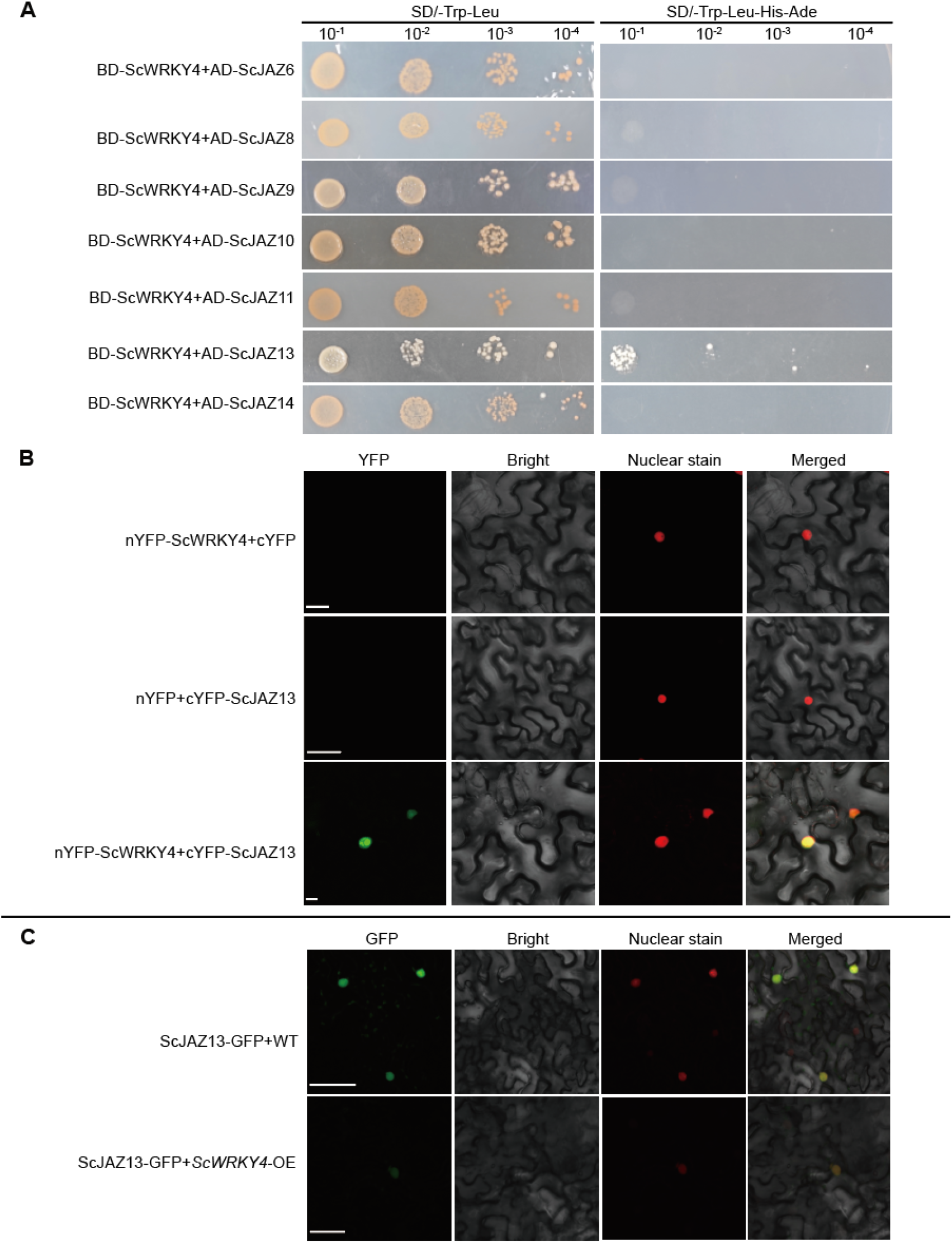
Sugarcane ScWRKY4 interacted with ScJAZ13 and inhibited its expression. (A) ScWRKY4 interacted with seven ScJAZs in yeast. BD: pGBKT7 vector, AD: pGADT7 vector. (B) BiFC to detect the interaction between ScWRKY4 and ScJAZ13 in *N. benthamiana* epidermal cells. Images were obtained with a confocal microscope at 2 dpi. Bars, 25 μm. (C) ScWRKY4 inhibited the expression of ScJAZ13 in *N. benthamiana* epidermal cells. Images were obtained with a confocal microscope at 2 dpi. Bars, 25 μm. WT, wild type *N. benthamiana*; *ScWRKY4*-OE, transgenic plants overexpressing *ScWRKY4*.

## Discussion

WRKY is one of the largest families of transcription factor in plants (Eulgem *et al*., 2000). Its function has been successively reported in different plant species (Wani *et al*., 2021), however, there were few reports on the function of WRKY in sugarcane (Javed *et al*., 2022; Li *et al*., 2020; Liu, 2012; Wang *et al*., 2022a; Wang *et al*., 2018a). Here in our study, *ScWRKY4* belonged to the class IIc WRKY family member, and could be induced by the stress of SA and MeJA (Fig. 3A). After inoculation with *S. scitamineum*, the expression was up-regulated in smut-susceptible sugarcane cultivars (YZ03-103 and FN39) and down-regulated in smut-resistant cultivars (YZ01-1413 and YT96-86) (Fig. 3A). It is thus speculated that *ScWRKY4* negatively regulated the resistance of sugarcane to smut pathogen infection.

Previous studies have shown that plant *WRKY* gene could respond to infection caused by pathogenic fungi (Abbruscato *et al*., 2012; Li *et al*., 2006). Interesting but not surprising is that ROS levels in plants altered when they were infected by pathogens (Li *et al*., 2015). After being stressed by pathogenic bacteria, the expression of those genes related to ROS scavenging system-related genes such as *CAT* and *GST,* was closely related to plant disease resistance (Yan *et al*., 2015). *GhWRKY27a* inhibited the expression of *CAT* and *GST* genes and negatively regulated tobacco resistance to *Rhizoctonia solani* (Yan *et al*., 2015). HR is a defense response of plants to pathogens in host-parasite incompatibility (Klement and Goodman, 1967). Overexpression of *CaWRKY40* in tobacco altered the expression of HR-related genes (*NtHSR201, NtHSR203* and *NtHSR515*) and pathogenicity-related genes, thereby regulating pepper response to *R. solanacearum* stress (Dang *et al*., 2013). In this study, the CAT enzyme activity and GSTs enzyme activity of OE were significantly higher than those of WT before inoculation, but decreased to the control level after inoculation (Fig. 4C). Besides, the expression of HR marker genes *NbHSR201* and *NbHSR515*, and the ROS-related genes *NbCAT1*, was significantly decreased (Fig. 4D, G). It is speculated that *N. benthamiana* plants overexpressing *ScWRKY4* may reduce tolerance to oxidative stress after inoculation with *F. solani* var. *coeruleum*. Reasobably, the inhibition of HR occurrence was associated with the decreased enzymatic activities of CAT and GSTs and the down-regulated expression levels of ROS and HR-related genes. SA and JA are two widely studied signaling pathways that play important regulatory roles in plant defense responses to pathogenic infection (Peng *et al*., 2012; Spoel and Dong, 2008). WRKY25 was a negative regulator of SA-mediated defense responses against *Pseudomonas syringae* in Arabidopsis WRKY25 T-DNA insertion mutants and transgenic plants overexpressing (Zheng *et al*., 2007). In rice, JA played an important role in *OsWRKY30* gene mediated defense responses to fungal pathogens (Peng *et al*., 2012). Here, the contents of SA and JA in OE were lower than those in the control at 2 dpi (Fig. 4B), and genes related to SA and JA signaling pathways (*NbPR2, NbPR3, NbLOX1* and *NbDEF1*) were down-regulated in OE (Fig. 4E, F). Therefore, it is assumed that ScWRKY4 is a negative regulatory transcription factor, which may suppress the SA and JA signaling pathways by inhibiting the expression of *NbPR2, NbPR3, NbLOX1* and *NbDEF1* genes, and at the same time repress the expression of HR and ROS-related genes, *NbHSR201, NbHSR515* and *NbCAT1*, thereby weakening the resistance of *N. benthamiana* to *F. solani* var. *coeruleum*.

What’s exciting is that plant *WRKY* genes are involved in different defense signaling pathways (Birkenbihl *et al*., 2012; Peng *et al*., 2012). Overexpression of rice *OsWRKY03* enhanced the resistance of transgenic plants to bacterial blight and induced the expression of several pathogenic related genes in transgenic plants (Liu *et al*., 2005). Further studies revealed that *OsWRKY03*, located upstream of *OsNPR1*, acts as a transcriptional activator of SA-related or JA-related defense signaling pathways (Liu *et al*., 2005). Besides, AtWRKY57 competed with AtWRKY33 to interact with VQ proteins SIB1 and SIB2, and competitively regulated the expression of key repressors JAZ1 and JAZ5 of the JA signaling pathway, thereby blocking JA signaling to a certain extent and attenuating the effect of WRKY33 on *Botrytis cinerea* resistance (Jiang and Yu, 2016). OsWRKY13 enhanced rice defense responses against *R. solani* and *Sarocladium oryzae*, which may affect the TIFY9-mediated MAPK cascade signaling pathway (Lilly and Subramanian, 2019). In the present study, transcriptome data showed that in the plant hormone signal transduction pathway, after inoculation of *F. solani* var. *coeruleum*, the expression of most *JAZ* genes was significantly suppressed in *N. benthamiana* plants overexpressing the *ScWRKY4* gene (Fig. 6A, Supplementary Table S8). We also confirmed that ScWRKY4 interacted with ScJAZ13 and could repress the expression of ScJAZ13 (Fig. 7A–C). Therefore, we reasonably deduced that ScWRKY4 may negatively regulate resistance to pathogens by repressing the expression of ScJAZ13.

In conclusion, we depicted here a model for the disease resistance regulatory mechanism of the *ScWRKY4* gene (Fig. 8). That is, stable overexpression of *ScWRKY4* negatively regulated the resistance of transgenic plants to pathogen infection and caused the down-regulated expression of JA-related genes. During the transcription process, most of the JAZ genes could be repressively expressed in *ScWRKY4* transgenic plants. In addition, ScJAZ13 protein interacted with ScWRKY4 protein and was repressed by ScWRKY4. We thus proposed that ScWRKY4 may enhance susceptibility to pathogen by repressing the expression of ScJAZ13. This work is expected to lay the foundation for in-depth analysis of the biological function and mechanism of sugarcane WRKY transcription factors.

**Fig. 8.**
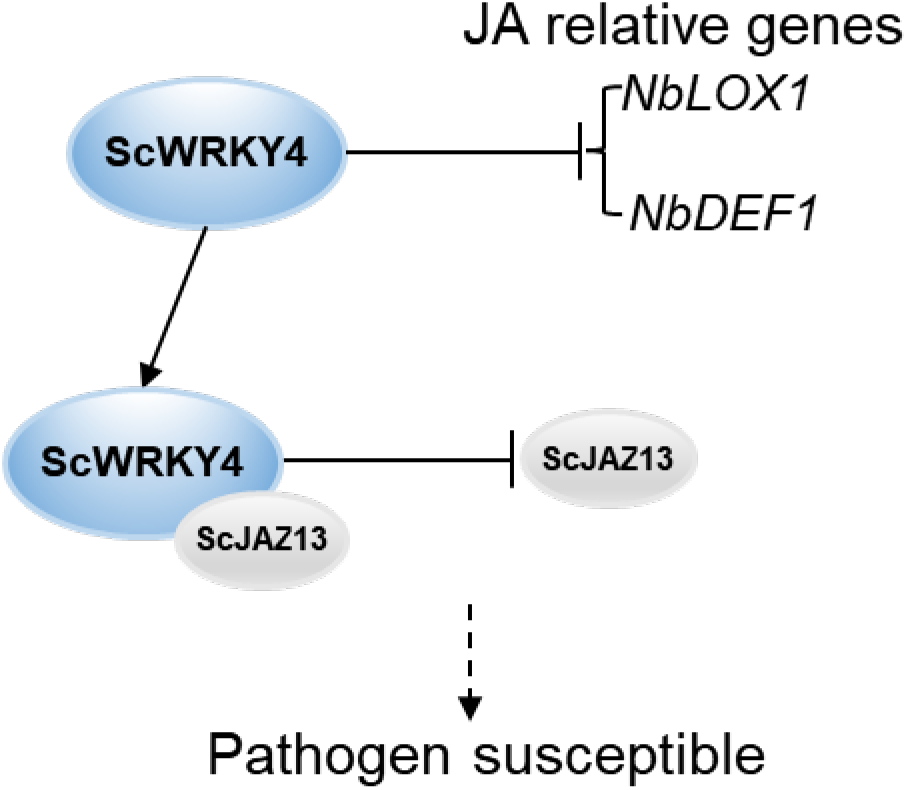
A model for disease resistance regulation of the transcription factor ScWRKY4. Stable overexpression of *ScWRKY4* caused down-regulated expression of JAZ-related genes in transgenic plants. ScWRKY4 interacted with ScJAZ13 and repressed the expression of ScJAZ13. Thus, it is hypothesized that ScWRKY4 enhances susceptibility to pathogen and is a negatively regulated transcription factor.

## Abbreviations

SA: salicylic acid
MeJA: methyl jasmonate
ROS: reactive oxygen species
JA: jasmonic acid
RT: qPCR-real-time quantitative PCR
DAB: 3,3’-diaminobenzidine
GO: gene ontology
BiFC: bimolecular fluorescent complimentary
YFP: yellow fluorescent protein
GFP: green fluorescent protein
WT: wild-type
GAPDH: glyceraldehyde-3-phosphate dehydrogenase
HR: hypersensitive response
ET: ethylene
DEGs: differentially expressed genes

## Supplementary Data

**Table S1.** Primers used in the experiment.

**Table S2.** GO annotation of ShWRKY gene family in *Saccharum*. spp. hybrid cultivar R570.

**Table S3.** The percentage between 60 ShWRKYs and ScWRKY4.

**Table S4.** Function prediction of *cis*-acting regulatory element of in the promoters of sugarcane ScWRKY4.

**Table S5.** The statistics information of RNA-seq alignment.

**Table S6.** The 30 most significant GO enrichment term for differentially expressed genes specific to the WT-CK-vs-WT-T.

**Table S7.** The 30 most significant GO enrichment term for differentially expressed genes specific to the *ScWRKY4*-CK-vs-*ScWRKY4*.

**Table. S8.** Differential gene expression in disease resistance related pathways.

**Fig. S1.** Screening of the T_3_ generation of transgenic *N. benthamiana* plants overexpressing *ScWRKY4*.

**Fig. S2.** The analysis on the RNA-seq data.

**Fig. S3.** The differentially expressed genes in WT and *ScWRKY4* gene overexpressing tobacco plants after inoculated with *F. solani* var. *coeruleum* at 0 dpi and 2 dpi.

## Author contributions

DJW and YXQ conceived and designed the experiments. DJW, WW, SJZ, LQQ, YLL, and PXL performed the experiments. DJW analyzed the data and wrote the paper. YCS and YXQ revised the final version of the paper. All authors read and approved the final manuscript.

## References

Abbruscato P, Nepusz T, Mizzi L, Del Corvo M, Morandini P, Fumasoni I, Michel C, Paccanaro A, Guiderdoni E, Schaffrath U, Morel JB, Piffanelli P, Faivre-Rampant O. 2012. *OsWRKY22*, a monocot *WRKY* gene, plays a role in the resistance response to blast. Molecular Plant Pathology 13, 828–841.

Bailey TL, Boden M, Buske FA, Frith M, Grant CE, Clementi L, Ren J, Li WW, Noble WS. 2009. MEME SUITE: tools for motif discovery and searching. Nucleic Acids Research 37, W202–208.

Bakshi M, Oelmüller R. 2014. WRKY transcription factors. Plant Signaling & Behavior 9, e27700.

Bhuiyan SA, Magarey RC, McNeil MD, Aitken KS. 2021. Sugarcane smut, caused by *Sporisorium scitamineum*, a major disease of sugarcane: a contemporary review. Phytopathology 111, 1905–1917.

Birkenbihl RP, Diezel C, Somssich IE. 2012. Arabidopsis WRKY33 is a key transcriptional regulator of hormonal and metabolic responses toward *Botrytis cinerea* infection. Plant Physiology 159, 266–285.

Cantalapiedra CP, Hernández-Plaza A, Letunic I, Bork P, Huerta-Cepas J. 2021. eggNOG-mapper v2: functional annotation, orthology assignments, and domain prediction at the metagenomic scale. Molecular Biology and Evolution 38, 5825–5829.

Chen CJ, Chen H, Zhang Y, Thomas HR, Frank MH, He YH, Xia R. 2020. TBtools: an integrative toolkit developed for interactive analyses of big biological data. Molecular Plant 13, 1194–1202.

Chen S, Zhou Y, Chen Y, Gu J. 2018. fastp: an ultra-fast all-in-one FASTQ preprocessor. Bioinformatics 34, i884–i890.

Connolly MA, Clausen PA, Lazar JG. 2006. Preparation of RNA from plant tissue using trizol. CSH Protocols 2006, pdb.prot4105.

Dang, FF, Wang, YN, Eulgem, Lai, Liu, ZQ, Qiu, AL. 2013. CaWRKY40, a WRKY protein of pepper, plays an important role in the regulation of tolerance to heat stress and resistance to *Ralstonia solanacearum* infection. Plant Cell Environment 36, 757–774.

Edgar RC. 2004. MUSCLE: multiple sequence alignment with high accuracy and high throughput. Nucleic Acids Research 32, 1792–1797.

Eulgem T, Rushton PJ, Robatzek S, Somssich IE. 2000. The WRKY superfamily of plant transcription factors. Trends in Plant Science 5, 199–206.

Garsmeur O, Droc G, Antonise R, Grimwood J, Potier B, Aitken K, Jenkins J, Martin G, Charron C, Hervouet C, Costet L, Yahiaoui N, Healey A, Sims D, Cherukuri Y, Sreedasyam A, Kilian A, Chan A, Van Sluys MA, Swaminathan K, Town C, Bergès H, Simmons B, Glaszmann JC, van der Vossen E, Henry R, Schmutz J, D’Hont A. 2018. A mosaic monoploid reference sequence for the highly complex genome of sugarcane. Nature Communications 9, 2638.

Gull A, Lone A, Wani N. 2019. “Biotic and abiotic stresses in plants.” Abiotic and Biotic Stress in Plants, n. pag.

Howe ES, Clemente TE, Bass HW. 2012. Maize histone H2B-mCherry: a new fluorescent chromatin marker for somatic and meiotic chromosome research. DNA Cell Biology 31, 925–938.

Javed T, Zhou JR, Li J, Hu ZT, Wang QN, Gao SJ. 2022. Identification and expression profiling of WRKY family genes in sugarcane in response to bacterial pathogen infection and nitrogen implantation dosage. Frontiers in Plant Science 13, 917953.

Jiang JJ, Ma SH, Ye NH, Jiang M, Cao JH, Zhang JH. 2017. WRKY transcription factors in plant responses to stresses. Journal Integrative Plant Biology 59, 86–101.

Jiang YJ, Yu DQ. 2016. WRKY57 regulates JAZ genes transcriptionally to compromise *Botrytis cinerea* resistance in *Arabidopsis thaliana*. Plant Physiology 171, 2771–2782.

Jiao ZG, Sun JL, Wang CQ, Dong YM, Xiao SH, Gao XL, Cao QW, Li LB, Li WD, Gao C. 2018. Genome-wide characterization, evolutionary analysis of *WRKY* genes in Cucurbitaceae species and assessment of its roles in resisting to powdery mildew disease. PLoS ONE 13, e0199851.

Klement Z, Goodman R. 1967. The hypersensitive reaction to infection by bacterial plant pathogens. Annual Review of Phytopathology 5, 17–44.

Langmead B, Salzberg SL. 2012. Fast gapped-read alignment with Bowtie 2. Nature Methods 9, 357–359.

Lescot M, Déhais P, Thijs G, Marchal K, Moreau Y, Van de Peer Y, Rouzé P, Rombauts S. 2002. PlantCARE, a database of plant *cis*-acting regulatory elements and a portal to tools for in *silico* analysis of promoter sequences. Nucleic Acids Research 30, 325–327.

Li J, Brader G, Kariola T, Tapio Palva E. 2006. WRKY70 modulates the selection of signaling pathways in plant defense. The Plant Journal 46, 477–491.

Li J, Brader G, Palva ET. 2004. The WRKY70 transcription factor: a node of convergence for jasmonate-mediated and salicylate-mediated signals in plant defense. Plant Cell 16, 319–331.

Li J, Wang J, Wang NX, Guo XQ, Gao Z. 2015. GhWRKY44, a WRKY transcription factor of cotton, mediates defense responses to pathogen infection in transgenic *Nicotiana benthamiana*. Plant Cell, Tissue and Organ Culture (PCTOC) 121, 127–140.

Li Z, Hua XT, Zhong WM, Yuan Y, Wang YJ, Wang ZC, Ming R, Zhang JS. 2020. Genome-wide identification and expression profile analysis of WRKY family genes in the autopolyploid *Saccharum spontaneum*. Plant Cell Physiology 61, 616–630.

Lilly JJ, Subramanian B. 2019. Gene network mediated by *WRKY13* to regulate resistance against sheath infecting fungi in rice (*Oryza sativa* L.). Plant Science 280, 269–282.

Liu JX, Que YX, Guo JL, Xu LP, Wu JY, Chen RK. 2012. Molecular cloning and expression analysis of a WRKY transcription factor in sugarcane. African Journal of Biotechnology 11, 6434–6444.

Liu XQ, Bai XQ, Qian Q, Wang XJ, Chen MS, Chu CC. 2005. OsWRKY03, a rice transcriptional activator that functions in defense signaling pathway upstream of OsNPR1. Cell Research 15, 593–603.

Livak KJ, Schmittgen TD. 2001. Analysis of relative gene expression data using real-time quantitative PCR and the 2(-Delta Delta C(T)) Method. Methods 25, 402–408.

Müller AJ, Mendel RR, Schiemann J, Simoens C, Inzé D. 1987. High meiotic stability of a foreign gene introduced into tobacco by *Agrobacterium*-mediated transformation. Molecular and General Genetics 207, 171–175.

Pandey SP, Somssich IE. 2009. The role of WRKY transcription factors in plant immunity. Plant Physiology 150, 1648– 1655.

Peng XX, Hu YJ, Tang XK, Zhou PL, Deng XB, Wang HH, Guo ZJ. 2012. Constitutive expression of rice *WRKY30* gene increases the endogenous jasmonic acid accumulation, *PR* gene expression and resistance to fungal pathogens in rice. Planta 236, 1485–1498.

Que YX, Su YC, Guo JL, Wu QB, Xu LP. 2014. A global view of transcriptome dynamics during *Sporisorium scitamineum* challenge in sugarcane by RNA-Seq. PLoS One 9, e106476.

Que YX, Xu LP, Xu JS, Zhang JS, Zhang MQ, Chen RK. 2009. Selection of control genes in Real-time qPCR analysis of gene expression in sugarcane. Chinese Journal Of Tropical Crops 030, 274–278.

Rajput MA, Rajput NA, Syed RN, Lodhi AM, Que YX. 2021. Sugarcane smut: current knowledge and the way forward for management. Journal of Fungi 7, 1095.

Rushton PJ, Somssich IE, Ringler P, Shen QJ. 2010. WRKY transcription factors. Trends in Plant Science 15, 247– 258.

Singels A, Jackson P, Inman-Bamber G. 2021. Chapter 21-Sugarcane. In: Sadras VO, Calderini DF, eds. Crop Physiology Case Histories for Major Crops: Academic Press, 674–713.

Singh KB, Foley RC, Oñate-Sánchez L. 2002. Transcription factors in plant defense and stress responses. Current Opinion in Plant Biology 5, 430–436.

Spoel SH, Dong X. 2008. Making sense of hormone crosstalk during plant immune responses. Cell host & microbe 3, 348–351.

Su WH, Ren YJ, Wang DJ, Su YC, Feng JF, Zhang C, Tang HC, Xu LP, Muhammad K, Que YX. 2020. The alcohol dehydrogenase gene family in sugarcane and its involvement in cold stress regulation. BMC Genomics 21, 521.

Su WH, Zhang C, Wang DJ, Ren YJ, Zhang J, Zang SJ, Zou WH, Su YC, You CH, Xu LP, Que YX. 2022. A comprehensive survey of the aldehyde dehydrogenase gene superfamily in *Saccharum* and the role of *ScALDH2B-1* in the stress response. Environmental and Experimental Botany 194, 104725.

Subramanian B, Gao S, Lercher MJ, Hu S, Chen WH. 2019. Evolview v3: a webserver for visualization, annotation, and management of phylogenetic trees. Nucleic Acids Research 47, W270–W275.

Sun TT, Chen Y, Feng AY, Zou WH, Wang DJ, Lin PX, Chen YL, You CH, Que YX, Su YC. 2023. The allene oxide synthase gene family in sugarcane and its involvement in disease resistance. Industrial Crops and Products 192, 116136.

Wang DJ, Wang L, Su WH, Ren YJ, You CH, Zhang C, Que YX, Su YC. 2020. A class III WRKY transcription factor in sugarcane was involved in biotic and abiotic stress responses. Scientific Reports 10, 20964.

Wang DJ, Zhang C, Que BB, You CH, Luo J, Su YC. 2022a. Identification of WRKY gene family in sugarcane cultivar and expression analysis under *Sporisorium scitamineum* stress. Journal of Fujian Agriculture and Forestry University (Natural Science Edition) 51, 14.

Wang L, Liu F, Dai M, Sun T, Su W, Wang C, Zhang X, Mao H, Su Y, Que Y. 2018a. Cloning and Expression Characteristic Analysis of *ScWRKY4* Gene in Sugarcane. Acta Agronomica Sinica 44, 13.

Wang L, Liu F, Zhang X, Wang WJ, Sun TT, Chen YF, Dai MJ, Yu SX, Xu LP, Su Y, Que YX. 2018b. Expression characteristics and functional analysis of the *ScWRKY3* gene from Sugarcane. International Journal of Molecular Sciences 19, 4059.

Wang N, Fan X, He MY, Hu ZY, Tang CL, Zhang S, Lin DX, Gan PF, Wang JF, Huang XL, Gao CX, Kang ZS, Wang XJ. 2022b. Transcriptional repression of *TaNOX10* by TaWRKY19 compromises ROS generation and enhances wheat susceptibility to stripe rust. Plant Cell 34, 1784–1803.

Wani SH, Anand S, Singh B, Bohra A, Joshi R. 2021. WRKY transcription factors and plant defense responses: latest discoveries and future prospects. Plant Cell Reports 40, 1071–1085.

Xie WY, Ke YG, Cao JB, Wang SP, Yuan M. 2021. Knock out of transcription factor *WRKY53* thickens sclerenchyma cell walls, confers bacterial blight resistance. Plant physiology 187, 1746–1761.

Xiong XP, Sun SC, Li YJ, Zhang XY, Sun J, Xue F. 2019. The cotton WRKY transcription factor *GhWRKY70* negatively regulates the defense response against *Verticillium dahliae*. The Crop Journal 7, 393–402.

Yan Y, Jia HH, Wang F, Wang C, Liu SC, Guo XQ. 2015. Overexpression of *GhWRKY27a* reduces tolerance to drought stress and resistance to *Rhizoctonia solani* infection in transgenic *Nicotiana benthamiana*. Frontiers in physiology 6, 265.

Yin WC, Wang XH, Liu H, Wang Y, Nocker S, Tu MX, Fang JG, Guo JQ, Li Z, Wang XP. 2022. Overexpression of *VqWRKY31* enhances powdery mildew resistance in grapevine by promoting salicylic acid signaling and specific metabolite synthesis. Horticulture Research 9, uhab064.

Zhang D, Gao F, Jakovlić I, Zou H, Zhang J, Li WX, Wang GT. 2020. PhyloSuite: An integrated and scalable desktop platform for streamlined molecular sequence data management and evolutionary phylogenetics studies. Molecular Ecology Resources 20, 348–355.

Zhang YJ, Wang LG. 2005. The WRKY transcription factor superfamily: its origin in eukaryotes and expansion in plants. BMC Evolutionary Biology 5, 1.

Zheng ZY, Mosher SL, Fan BF, Klessig DF, Chen ZX. 2007. Functional analysis of *Arabidopsis* WRKY25 transcription factor in plant defense against *Pseudomonas syringae*. BMC Plant Biology 7, 2.

Zhou R, Dong YH, Liu X, Feng S, Wang CG, Ma XM, Liu JN, Liang Q, Bao Y, Xu SG, Lang XY, Gai SS, Yang KQ, Fang HC. 2022. JrWRKY21 interacts with JrPTI5L to activate the expression of *JrPR5L* for resistance to *Colletotrichum gloeosporioides* in walnut. Plant Journal 111, 1152–1166.

